# NOCICEPTOR NEURONS CONTROL POLLUTION-MEDIATED NEUTROPHILIC ASTHMA

**DOI:** 10.1101/2024.08.22.609202

**Authors:** Jo-Chiao Wang, Amelia Kulle, Theo Crosson, Amin Reza Nikpoor, Surbhi Gupta, Anais Roger, Moutih Rafei, Ajitha Thanabalasuriar, Sebastien Talbot

## Abstract

The immune and sensory nervous systems, having evolved in parallel, communicate through shared receptors and transmitters to maintain homeostasis and respond to both external and internal disruptions. Although neural responses often confer protective benefits, they can also exacerbate inflammation during allergic reactions such as asthma. In our study, we modeled pollution-exacerbated asthma by exposing mice to ambient PM_2.5_ particles alongside ovalbumin. Compared to exposure to ovalbumin alone, this co-exposure significantly increased the numbers of neutrophils and γδ T cells in bronchoalveolar lavage fluid. We found that silencing nociceptor neurons at the peak of inflammation using intranasal QX-314 or ablating TRPV1-expressing neurons reduced lung neutrophil accumulation. Live *in vivo* intravital imaging confirmed that neuronal ablation reduced neutrophil numbers and increased their net displacement capacity. In neurons isolated from mice with pollution-exacerbated asthma, the chemical-sensing TRPA1 channel exhibited heightened sensitivity to its cognate ligand. Elevated levels of artemin were detected in the bronchoalveolar lavage fluid of pollution-exposed mice but returned to baseline in mice with ablated nociceptor neurons. Alveolar macrophages expressing the pollution-sensing aryl hydrocarbon receptor were identified as a putative source of artemin following exposure to PM_2.5_. This molecule enhanced TRPA1 responsiveness and, in turn, drove nociceptor-mediated neutrophil recruitment, revealing a novel mechanism by which lung-innervating neurons respond to air pollution in the context of allergy. Overall, our findings suggest that targeting artemin-driven pathways could provide a therapeutic strategy for controlling neutrophilic airway inflammation in asthma, a clinical condition typically refractory to treatment.

## INTRODUCTION

Organisms have evolved sophisticated fail-safe systems to preserve homeostasis, integrating threat detection, reflex responses, and tailored immune mechanisms^1^. These systems are jointly orchestrated by the immune and sensory nervous systems, which are designed to sense external threats and internal disturbances. Both systems rely on shared metabolic pathways and signaling molecules—including receptors, cytokines, and neuropeptides ^2^. This common molecular framework supports constant interactions between these systems, which play pivotal roles in monitoring barrier tissues (e.g., skin ^3^, lung^4,5^, and gut^6^), activating anticipatory immune responses^7^, and managing pathologies such as allergies^8,9^, infections^10^, and malignancies^11–14^.

In the context of lung disease, vasoactive intestinal peptide (**VIP**), released by pulmonary sensory neurons following IL-5 stimulation, fosters allergic inflammation by acting on CD4⁺ T cells and innate lymphoid type 2 cells (**ILC2s**). This action elevates T helper type 2 (**T_h_2**) cytokines, which are central drivers of asthmatic responses^15,16^. Expanding on these insights, our work shows that vagal nociceptor neurons contribute to airway hyperresponsiveness^1,17^ mucus metaplasia^18,19^, and the detection of immunoglobulins E and G^20,21^. These neurons also mediate antibody class switching in B cells^22^ and promote IgG production^23^. More recently, we discovered that a subset of nociceptors is reprogrammed by the asthma-associated cytokine IL-13, rendering them susceptible to the inhibitory influence of neuropeptide Y (**NPY**) secreted by lung-innervating sympathetic neurons^24^.

Wildfire frequency and severity increased by 2.2-fold within the last 20 years ^25^. Between 2008 and 2018 in California, wildfire-attributable PM_2.5_ exposure caused an estimated ∼53,000 premature deaths^26^. Combined with urban pollution, these events have driven escalating levels of particulate matter (PM_2.5_), shifting asthma phenotypes from more responsive T helper 2 (T_h_2) / eosinophilic types to treatment-resistant neutrophilic and mixed T_h_17/T_h_1 variants^27–32^. Thus, recent estimates suggest that neutrophilic asthma accounts for 15–25% of all asthma cases, with about half of these resistant to standard therapies^33^.

To investigate whether neuronal silencing can mitigate neutrophilic airway inflammation, we used an innovative asthma model which combines ovalbumin with fine particulate matter (**FPM**). We further explored the molecular mechanisms underlying heightened neuronal sensitivity in this setting, identifying artemin—produced by alveolar macrophages via the pollution-sensing aryl hydrocarbon receptor (**AhR**)—as a critical mediator of this hypersensitivity.

## RESULTS

### Heightened nociceptor sensitivity in pollution-exacerbated asthma

Prof. Ya-Jen Chang’s group developed a novel method to model pollution-exacerbated asthma and investigate the impact of fine particulate matter (**FPM**) on airway inflammation ^34^. Their work demonstrates that FPM exposure induces airway hyperreactivity and neutrophilic inflammation, promotes T_h_1 and T_h_17 immune responses, and increases epithelial cell apoptosis rates ^34^. Notably, γδ T cells significantly contribute to both the inflammation and airway hyperreactivity phenotypes by producing IL-17A ^34^. Building on this foundation, we explored whether lung-innervating nociceptor neurons become sensitized during such asthma exacerbations. To test this, male and female C57BL/6 mice (6–10 weeks old) were sensitized with an intraperitoneal injection of ovalbumin (**OVA**; 200 µg/dose) and aluminum hydroxide (1 mg/dose) emulsion on days 0 and 7, followed by intranasal challenges of OVA (50 µg/dose), with or without FPM (20 µg/dose), on days 14–16. On day 17, lung-innervating jugular-nodose-complex (**JNC**) neurons were harvested, cultured for 24 hours, and loaded with the calcium indicator Fura-2AM (**Fig. 1A**). Stimulation with the TRPA1 agonist AITC (10–100 µM) and the pan-neuronal activator KCl (40 mM) revealed increased responsiveness in neurons from mice exposed to both FPM and OVA, compared to OVA-only–exposed mice, whose sensitivity was similar to that of wild-type controls (**Fig. 1B–C**).

**Figure 1.**
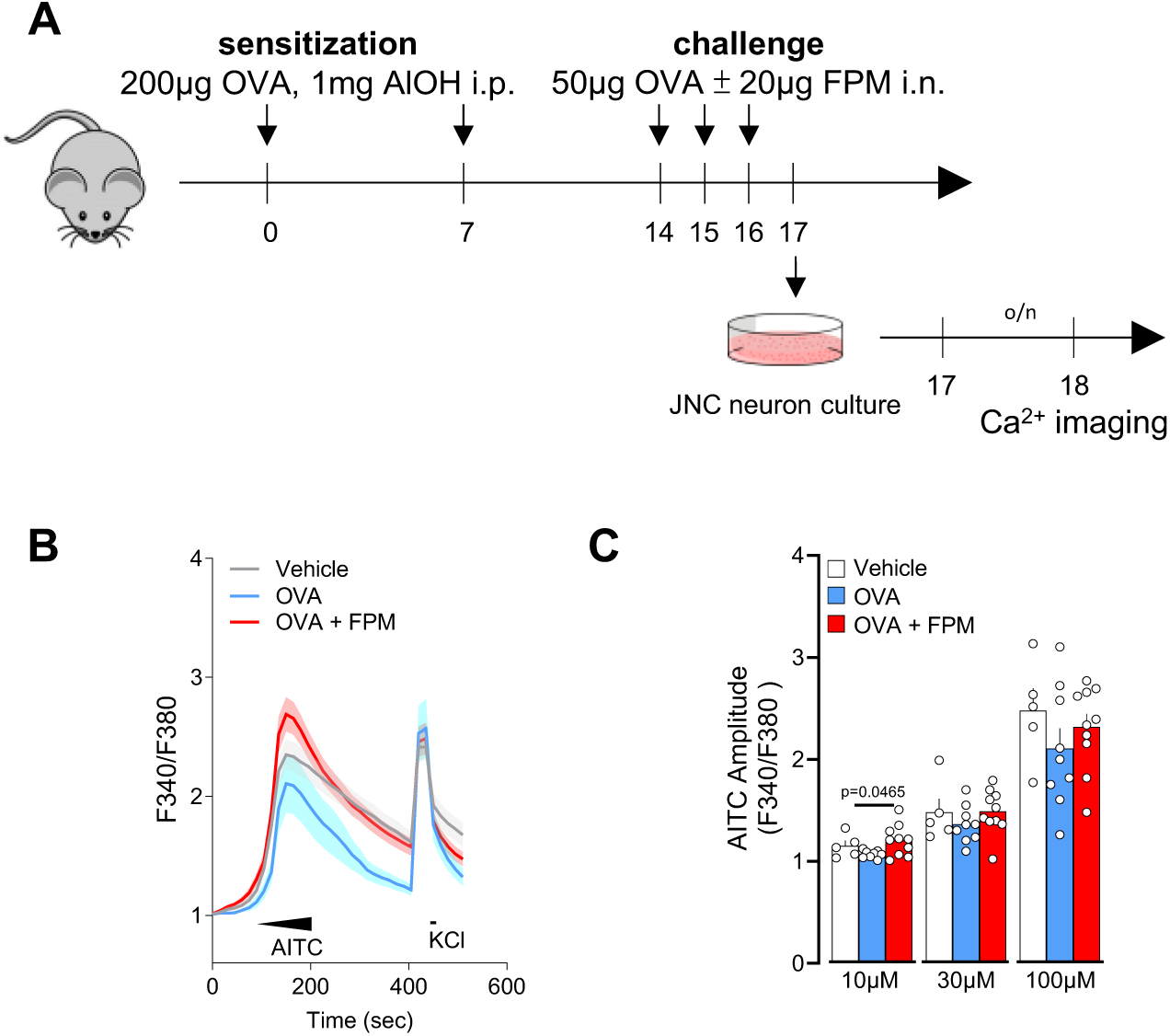
Air pollution exacerbates nociceptor neuronal activity. **(A–C)** Male and female C57BL/6 mice (6–10 weeks old) were sensitized via intraperitoneal injection with an emulsion of ovalbumin (OVA; 200 µg/dose) and aluminum hydroxide (1 mg/dose) on days 0 and 7. On days 14–16, mice were challenged intranasally with OVA (50 µg/dose), either alone or in combination with fine particulate matter (FPM; 20 µg/dose). Bronchoalveolar lavage fluid was collected, and jugular-nodose complex neurons were cultured on day 17 for 24 hours before being loaded with the calcium indicator Fura-2AM. Cells were sequentially stimulated with the TRPA1 agonist AITC (successively to 10 µM at 60–90 seconds, 30 µM at 90–120 seconds, 100 µM at 120–150 seconds) and then with KCl (40 mM at 420–435 seconds). Calcium flux was continuously monitored throughout the experiment. The amplitude of AITC responses was measured by calculating the ratio of peak F340/F380 fluorescence after stimulation to the baseline F340/F380 fluorescence measured 30 seconds prior to stimulation. Data are plots as the per dish average of AITC and KCl responsive neurons and show that AITC (10 µM) responses was higher in JNC neurons from OVA-FPM exposed mice when compared to vehicle or OVA alone (**C**). *Data in are presented as means ± SEM (**B-C**). N are as follows: **B**: n = 35 neurons (control group), 19 neurons (OVA group), and 38 neurons (OVA + FPM group), **C**: n = 5 dishes totalling 35 neurons (control group), 8 dishes totalling 42 neurons (OVA group), and 10 dishes totalling 76 neurons (OVA + FPM group). P-values were determined by nested one-way ANOVA with post-hoc Bonferroni’s. P-values are shown in the figure*.

### Transcriptomic reprogramming of TRPV1⁺ neurons

Recent data indicate that the asthma-associated cytokines IL-4 and IL-13 can reprogram lung-innervating nociceptor neurons to adopt a pro-allergic phenotype ^24^. To investigate whether pollution-exacerbated asthma similarly reprograms these neurons, we used TRPV1^cre^::tdTomato^fl/wt^ mice subjected to our standard OVA protocol, with or without FPM. We isolated and dissociated TRPV1⁺ JNC neurons, enriched them by excluding satellite glial and immune cells, and then purified them by FACS-sorting followed by RNA sequencing. Our analysis revealed that several differentially expressed genes (**DEGs**) were upregulated in the OVA-FPM group compared to naive mice (*Lifr, Oprm1*) (**Fig. 2A–B**), in OVA-FPM compared to OVA alone (*Oprm1, Nefh, P2ry1, Prkcb, Gabra1, Kcnv1*) (**Fig. 2C–D**), and in OVA alone compared to naive controls (*Cntn1, Piezo1, Npy1r, Kcna1*) (**Fig. 2E–F**). Collectively, these results suggest that pollution-exacerbated asthma both transcriptomically and functionally reprograms lung-innervating nociceptor neurons (**Supplementary Table 1**).

**Figure 2.**
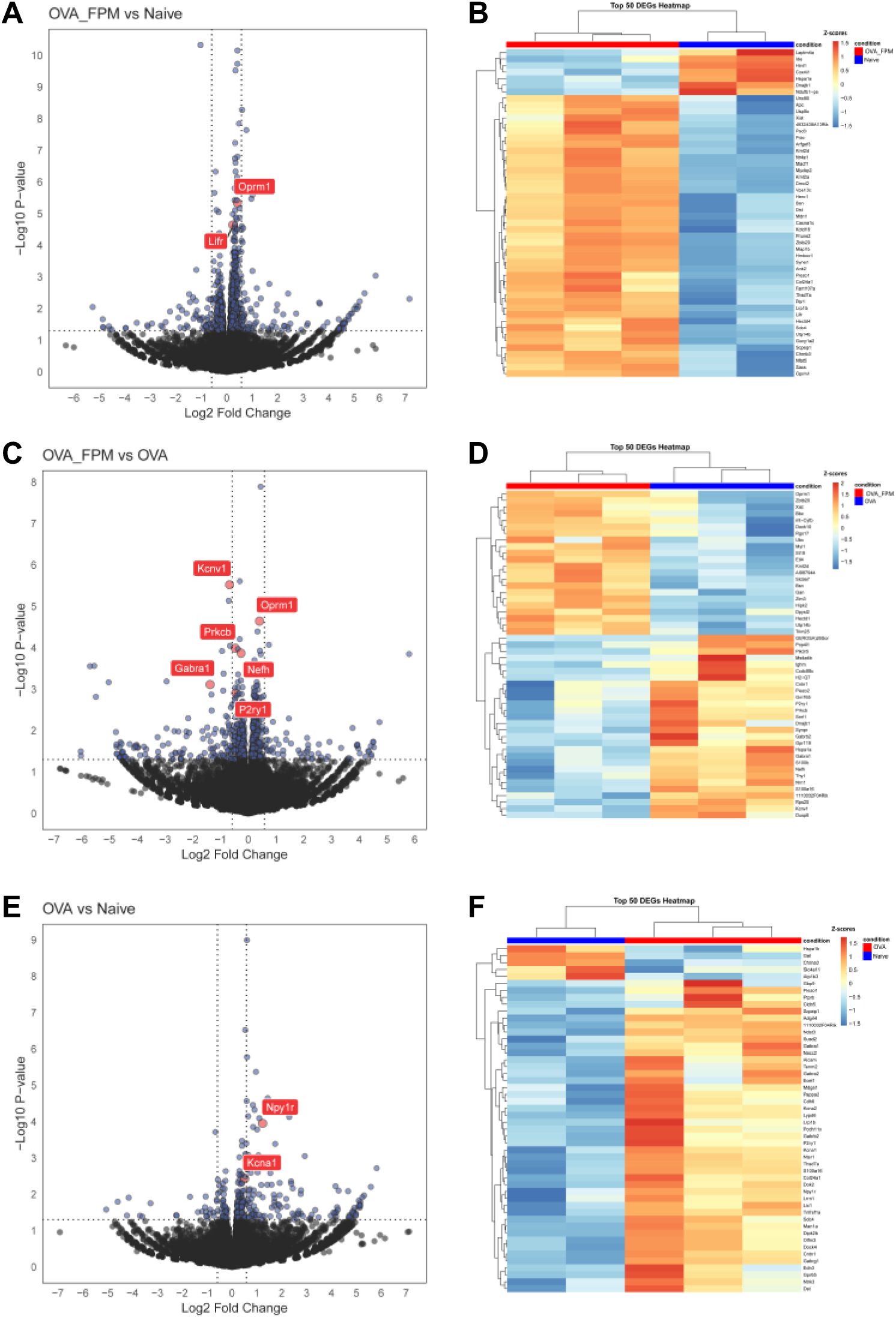
Air pollution reprograms the transcriptome of nociceptor neurons. (**A–F**) Naïve male and female TRPV1^cre^::tdTomato^fl/wt^ mice (6–10 weeks old) underwent either a pollution-exacerbated asthma protocol, the classic OVA protocol, or remained naïve. On day 17 (peak inflammation), jugular-nodose-complex (**JNC**) neurons were harvested and dissociated, and TRPV1⁺ neurons (tdTomato⁺) were sorted via FACS to remove stromal cells and non-peptidergic neurons. RNA was then isolated for sequencing. Volcano plots (**A, C, E**) and heatmaps (**B, D, F**) show differentially expressed genes (**DEGs**) for three comparisons: OVA + FPM vs. naïve (**A-B**), OVA + FPM vs. OVA alone (**C-D**), and OVA alone vs. naïve (**E-F**). Notable genes with increased expression include *Lifr* and *Oprm3* in OVA + FPM vs. naïve, *Oprm1, Nefh, P2ry1, Prkcb, Gabra1,* and *Kcnv1* in OVA + FPM vs. OVA, and *Npy1r* and *Kcna1* in OVA alone vs. naïve. *Data are presented either as volcano plots (**A, C, E**), showing the log2 fold change of TPM between groups along with the corresponding –log10 p-values from DESeq2 analysis, or as heatmaps (**B, D, F**), showing the z-scores of rlog-transformed normalized counts. The experimental groups were naïve (n=2; **A–B, E–F**), OVA (n=3; **C–F**), and OVA-FPM (n=3; **A–D**). P-values were determined by DESeq2 (**A, C, E**) and are indicated in the figure*.

### Silencing nociceptor neurons reduces inflammation

To determine whether silencing nociceptor neurons affects pollution-exacerbated asthma, we adapted a previously established neuron-blocking strategy originally developed for pain and itch neurons. This method uses nonselective ion channels (TRPA1 and TRPV1) as a drug entry port for QX-314, a charged form of lidocaine. During inflammation, QX-314 enters sensory fibers, blocking sodium currents and producing a targeted, long-lasting (>9 hours) electrical blockade of nociceptors without impairing immune cell function^35–38^. Intranasal administration of QX-314 has already been shown to reverse allergic airway inflammation, coughing, mucus metaplasia, and hyperreactivity^16,36,39–41^.

In our model, an intranasal dose of QX-314 administered at the peak of inflammation (day 16, 5 nmol, 50 µL) significantly reduced neutrophil counts in bronchoalveolar lavage fluid (**BALF**) of OVA-FPM-exposed mice, restoring them to levels typically seen in asthmatic (OVA-only) mice (**Fig. 3A–B**). To validate these findings, we used mice genetically engineered to either retain (TRPV1^wt^::DTA^fl/wt^; denoted as TRPV1^WT^) or lack (TRPV1^cre^::DTA^fl/wt^; denoted as TRPV1^DTA^) TRPV1-expressing nociceptors^11,42,43^. Compared with controls, TRPV1^DTA^ mice had markedly fewer BALF neutrophils and lung γδ T cells (**Fig. 3C–E**), underscoring the critical role nociceptor neurons play in shaping the inflammatory and immune responses in pollution-exacerbated asthma.

**Figure 3.**
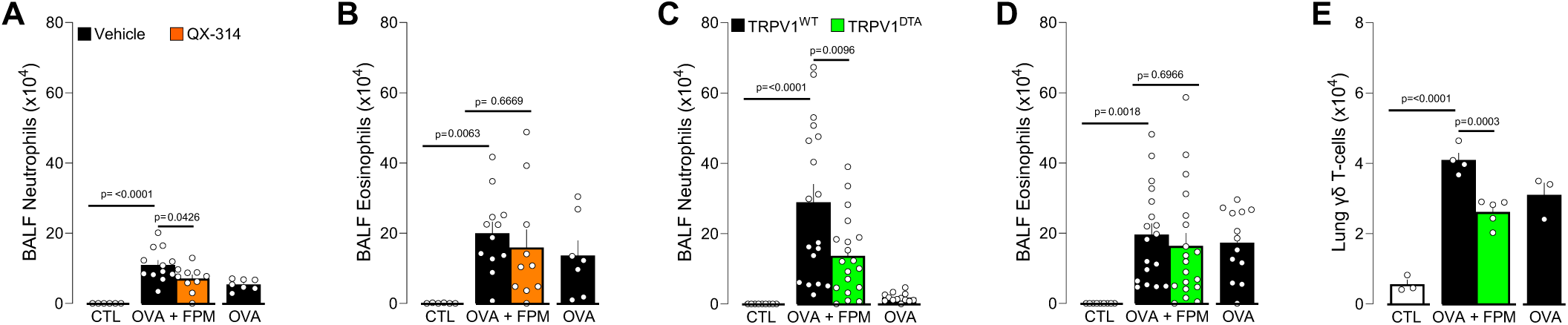
Nociceptor neurons control pollution-exacerbated asthma. **(A–B)** Male and female C57BL/6 mice (6–10 weeks old) were sensitized intraperitoneally with ovalbumin (OVA; 200 µg/dose in 200 µL) and aluminum hydroxide (1 mg/dose in 200 µL) on days 0 and 7. On days 14–16, mice were challenged intranasally with OVA (50 µg/dose in 50 µL) alone or with fine particulate matter (FPM; 20 µg/dose in 50 µL). On day 16, 30 minutes after the final challenge, mice received intranasal QX-314 (5 nmol/dose in 50 µL). Bronchoalveolar lavage fluid (BALF) was collected on day 17 and analyzed by flow cytometry. Compared with naïve or OVA-exposed mice, those co-challenged with OVA + FPM showed increased BALF neutrophils (**A**). QX-314 treatment normalized these levels, while BALF eosinophil levels remained comparable (**B**). **(C–E)** Male and female littermate control (TRPV1^WT^) and nociceptor-ablated (TRPV1^DTA^) mice (6–10 weeks old) were sensitized and challenged under the same OVA ± FPM protocol (days 0, 7, and 14–16). BALF or lungs were collected on day 17 and assessed by flow cytometry. Compared with naïve or OVA-exposed mice, OVA + FPM co-challenged mice exhibited higher BALF neutrophils (**C**) and lung γδ T cells (**E**). Nociceptor ablation protected against these increases (**C, E**), while BALF eosinophil levels remained comparable (**D**). *Data are shown as mean ± SEM (**A–E**). Experiments were replicated twice, and animals pooled (**A-E**). N are as follows: **A-B**: control (n=6), OVA (n=7), OVA-FPM (n=12), OVA-FPM+QX-314 (n=10), **C-D**: TRPV1^WT^ + control (n=9), TRPV1^WT^ + OVA (n=13), TRPV1^WT^ + OVA-FPM (n=18), TRPV1^DTA^ + OVA-FPM (n=19), **E**: TRPV1^WT^ + control (n=3), TRPV1^WT^ + OVA (n=3), TRPV1^WT^ + OVA-FPM (n=4), TRPV1^DTA^ + OVA-FPM (n=5). P-values were determined by a one-way ANOVA with post-hoc Tukey’s (**A-E**). P-values are shown in the figure*.

### Impact on myeloid cell motility

Building on these observations, we used intravital microscopy^44,45^ to understand how pollution-exacerbated asthma and lung innervation influence the motility of alveolar macrophages (**AMs**) and neutrophils. Although the total number of AMs (labelled by i.n. PKH26) remained unchanged (**Fig. 4A-B**), their net displacement was lower in nociceptor-ablated (Na_V_1.8^cre^::DTA^fl/wt^; denoted as Na_V_1.8^DTA^) mice than in littermate controls (Na_V_1.8^wt^::DTA^fl/wt^; denoted as Na_V_1.8^WT^; **Fig. 4C-F, Supplementary Video 1**). In contrast, neutrophil accumulation (labelled by i.v. αLy6G) was observed in pollution-exposed, asthmatic littermate controls, but was absent in nociceptor-ablated mice (**Fig. 4G-H**). These data are consistent with our BALF flow cytometry data (**Fig. 3A, 3C**). Furthermore, total neutrophil displacement was higher in Na_V_1.8^DTA^ mice (**Fig. 4I–L, Supplementary Video 2**), without any bias toward a specific behavior subtype (e.g. adherent, crawling, patrolling, or tethering; **Fig. 4L**). Taken together, these data suggest that nociceptor neurons modulate AM motility and neutrophil accumulation while limiting the motility of recruited neutrophils.

**Figure 4.**
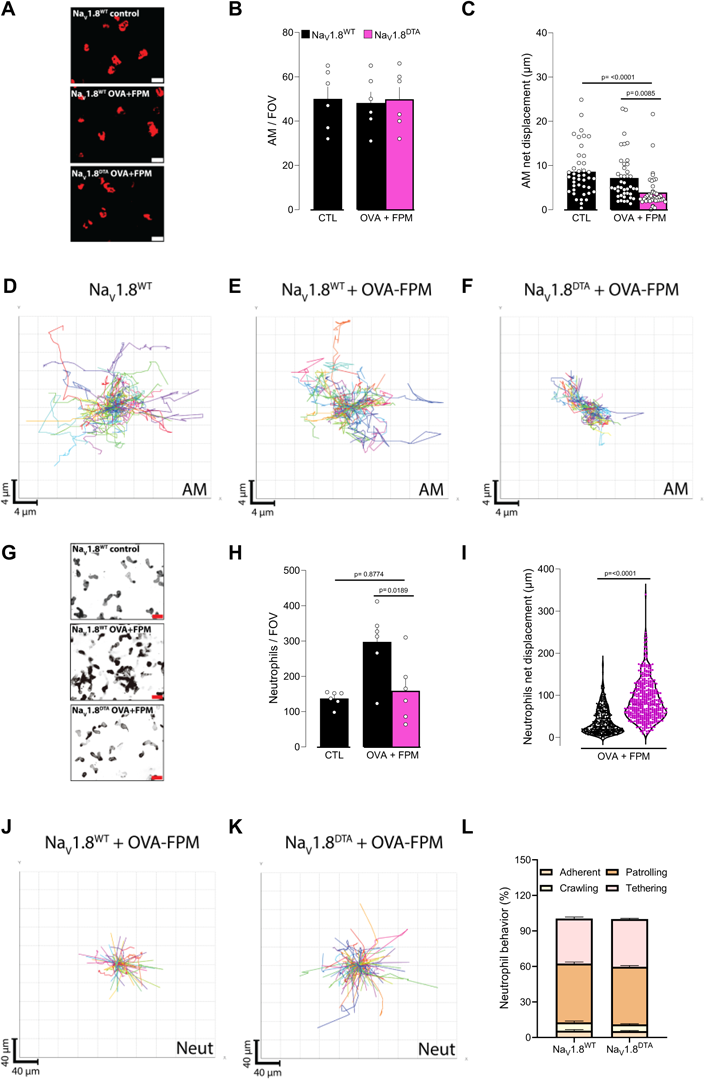
Vagal sensory neurons gatekeep alveolar macrophage motility and neutrophil numbers. **(A–D)** Male and female littermate control (Na_V_1.8^wt^::DTA^fl/wt^ denoted as Na_V_1.8^WT^) and nociceptor-ablated (Nav1.8^cre^::DTA^fl/wt^ denoted as NaV1.8^DTA^) mice (6–10 weeks old) were sensitized via intraperitoneal injection with an emulsion of ovalbumin (OVA; 200 µg/dose) and aluminum hydroxide (1 mg/dose) on days 0 and 7. Phagocytes were labeled by intranasal injection of PKH26 Red Fluorescent Cell Linker Kit (25 pmol/dose) on day 10. Mice were then challenged intranasally with OVA (50 µg/dose) alone or in combination with fine particulate matter (FPM; 20 µg/dose) on days 14–16, and images were acquired on day 17. **(A)** Representative maximum-intensity projection of alveolar macrophages (AMs, red). Scale bar: 20 µm. **(B)** Quantification of AM numbers per field of view (FOV). **(C)** Net displacement of AMs over 1 hour. **(D-F)** Representative tracks of individual AMs (each color represents a single AM) over 1 hour. While the AM numbers (**A-B**) were not impacted, their net displacement (**C-F**) was reduced in OVA-FPM-exposed NaV1.8^DTA^ mice. **(G-L)** Male and female littermate control (Na_V_1.8^wt^::DTA^fl/wt^ denoted as Na_V_1.8^WT^) and nociceptor-ablated (Na_V_1.8^cre^::DTA^fl/wt^ denoted as Na_V_1.8^DTA^) mice (6–10 weeks old) were sensitized via intraperitoneal injection with an emulsion of ovalbumin (OVA; 200 µg/dose) and aluminum hydroxide (1 mg/dose) on days 0 and 7. Mice were then challenged intranasally with OVA (50 µg/dose) alone or in combination with fine particulate matter (FPM; 20 µg/dose) on days 14–16, and images were acquired on day 17. Immediately prior to perform the intravital imaging, we administered an intravenous Ly6G antibody to label neutrophils. **(G)** Representative maximum-intensity projection of neutrophils (black). Scale bar: 20 µm. **(H)** Quantification of neutrophil numbers per FOV. **(I)** The total displacement of neutrophils over 20 minutes. (**J-K)** Representative tracks of individual neutrophils (each color represents a single neutrophil) over 20 minutes. **(L)** Frequency of neutrophil behaviors per FOV (adherent, crawling, patrolling, or tethering). Data show that OVA-FPM-exposed littermate control mice present an increase in neutrophil numbers per field of view (**G-H**), an effect absent in Na_V_1.8^DTA^. Interestingly, OVA-FPM-exposed Na_V_1.8^DTA^ mice show an increase in neutrophil net displacement (**I-K**), an effect irrespective to one of the specific behaviors tested (**L**). *Data are presented as representative image (**A, G**; scale bar: 20µm), mean ± SEM (**B, C, H**), spider plot (**D-F, J-K**), violin plot showing median (**I**), and staked bar graph showing mean ± SEM (**L**). N are as follows: **B**: Na_V_1.8^WT^ + control (n=6), Na_V_1.8^WT^ + OVA-FPM (n=6), Na_V_1.8^DTA^ + OVA-FPM (n=6), **C**: Na_V_1.8^WT^ + control (n=42), Na_V_1.8^WT^ + OVA-FPM (n=42), Na_V_1.8^DTA^ + OVA-FPM (n=42), **H**: Na_V_1.8^WT^ + control (n=6), Na_V_1.8^WT^ + OVA-FPM (n=6), Na_V_1.8^DTA^ + OVA-FPM (n=6), **I**: Na_V_1.8^WT^ + OVA-FPM (n=385), Na_V_1.8^DTA^ + OVA-FPM (n=469), **J**: Na_V_1.8^WT^ + OVA-FPM (n=10), Na_V_1.8^DTA^ + OVA-FPM (n=11). P-values were determined by a one-way ANOVA with post-hoc Tukey’s (**B, C, H**) or unpaired Student T-test (**I, L**). P-values are shown in the figure*.

### Elevated cytokines and artemin under pollution-exacerbated asthma

To further explore the role of nociceptor neurons in airway inflammation, we performed an unbiased multiplex cytokine array. This revealed elevated levels of the asthma-promoting cytokine IL-4 and TNFα, with TNFα levels returning to baseline following nociceptor ablation (**Fig. 5A**). In addition, targeted ELISA showed increased artemin levels, which returned to normal in the absence of nociceptor neurons (**Fig. 5B**). Artemin, a member of the glial cell line-derived neurotrophic factor (**GDNF**) family, is pivotal for the development and function of sympathetic^46^ and sensory^47^ neurons through its interaction with the GFRα3-RET receptor complex. Activation of this receptor supports neuronal survival and growth during embryogenesis and drives neuronal hyperactivity under inflammatory conditions^48–50^.

**Figure 5.**
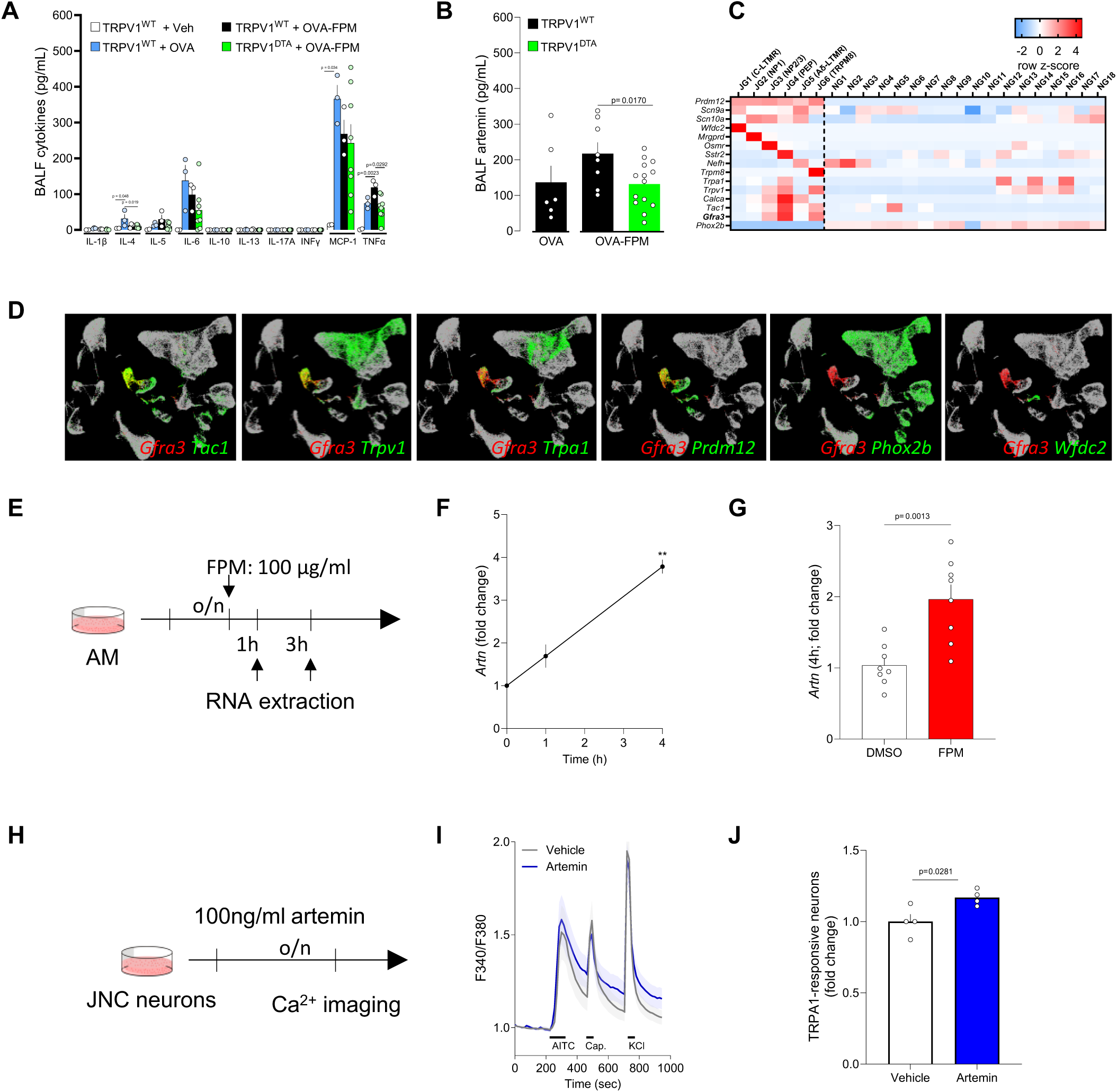
Artemin sensitizes TRPA1 activity in vagal sensory neurons. **(A-B)** Male and female littermate control (TRPV1^WT^) and nociceptor-ablated (TRPV1^DTA^) mice (6–10 weeks old) were sensitized and challenged under the same OVA ± FPM protocol (days 0, 7, and 14–16). BALF was collected on day 17 and assessed by multiplex array and ELISA. Compared with naïve or OVA-alone groups, OVA + FPM co-challenged mice exhibited levels of TNFα, and artemin. Notably, ablating nociceptors prevented these increases. **(C)** In-silico analysis of the GSE124312 dataset^51^. The heatmap displays transcript expression levels for the pan neural-crest lineage transcription factor (*Prdm12*), voltage-gated sodium channels (*Scn9a, Scn10a*), jugular subset markers (*Wfdc2, Mrgprd, Osmr, Sstr2, Nefh, Trpm8*), peptidergic neuron markers (*Trpa1, Trpv1, Calca, Tac1, Gfra3*), and the pan placodal lineage marker (*Phox2b*). Gfra3 expression is enriched in the peptidergic neuron cluster labeled JG4. Experimental details and cell clustering are described by Kupari et al^51^. **(D)** In-silico analysis of GSE192987^52^ showing co-expression of *Gfra3* with *Trpa1* and other inflammatory markers. Data are visualized as row z-scores in a heatmap or via UMAPs (TPTT > 1). Experimental details and cell clustering are described by Zhao et al^52^. **(E-G)** Alveolar macrophages (3 × 10⁵ cells/well) from naïve male and female C57BL/6 mice were cultured overnight and then stimulated with vehicle (DMSO) or FPM (100 µg/mL). RNA was extracted 1- and 4-hours post-stimulation and *Artn* expression was assessed using qPCR. FPM exposure increased *Artn* transcript levels at both 1 and 4 hours (**F, G**). **(H-J)** Naïve mice jugular-nodose-complex neurons were harvested, pooled, and cultured overnight with either vehicle or artemin (100 ng/mL). Cells were sequentially stimulated with AITC (TRPA1 agonist; 300 µM at 240–270 seconds), capsaicin (TRPV1 agonist; 300 nM at 320–335 seconds), and KCl (40 mM at 720–735 seconds). The percentage of AITC-responsive neurons (among all KCl-responsive cells) was normalized to vehicle-treated controls for each batch of experiments. Artemin-treated neurons showed increased responsiveness to AITC, while responses to capsaicin and KCl were unchanged (**I-J**). *Data are presented as means ± SEM (**A-B, F-G, J**), heatmap displaying the z-score of DESeq2 normalized counts **(C),** tSNE plots (**D**), schematics (**E, H**), means ± 95% CI of maximum Fura-2AM (F/F₀) fluorescence (**I**). N are as follows: **A**: TRPV1^WT^ + control (n=2), TRPV1^WT^ + OVA (n=3) TRPV1^WT^ + OVA-FPM (n=3), TRPV1^DTA^ + OVA-FPM (n=8), **B**: TRPV1^WT^ + OVA (n=6) TRPV1^WT^ + OVA-FPM (n=8), TRPV1^DTA^ + OVA-FPM (n=14), **F**: n=2/time point, **G**: n=8/group, **I**: vehicle (n=107 neurons), Artemin (n=122 neurons); **J**: n=4/group. P-values were determined by a one-way ANOVA with post-hoc Tukey’s (**A**, **B**) or unpaired Student T-test (**G, J**). P-values are shown in the figure*.

### Mechanistic insights: artemin–nociceptor axis

To determine which vagal neuron subsets, express the artemin receptor GFRα3–RET, we performed an in-silico analysis of single-cell RNA sequencing data^51,52^ and found that *Gfra3* is expressed by several nociceptor subtypes **(Fig. 5C–D)**, including JG3 (OSM-R–expressing), JG4 (TRPA1/SSTR2-expressing), and JG6 (TRPM8-expressing). Concurrently, ImmGen data^53^ indicated robust *Artn* expression in macrophages, with *Ahr* levels slightly lower than those in ILC2s and eosinophils **(SF. 1)**. We then conducted an in-silico analysis of data from Karlsson et al.^54^ **(SF. 2A)**, Uhlén et al. ^55^ **(SF. 2B)** and Abdulla et al. ^56^ **(SF. 2C)**, showing that *Ahr* is also expressed in human lung, that AHR protein is detected in human lung macrophages, and that *Artn* and *Ahr* are co-expressed in lung macrophages, respectively. Guided by these findings, we isolated alveolar macrophages from naive C57BL/6 mice and exposed them to fine particulate matter (FPM), observing a time-dependent increase in *Artn* transcripts **(Fig. 5E–G)**. These results highlight key cellular interactions involving FPM, AhR and artemin in the context of lung inflammation.

A recent study^49^ linked artemin overexpression to increased mRNA levels of several nociceptor markers (*Gfra3*, *TrkA*, *Trpv1*, *Trpa1*), accompanied by heightened thermal sensitivity. Consistent with this, our experiments in JNC neurons from OVA-FPM co-exposed mice revealed amplified calcium responses to the TRPA1 agonist AITC **(Fig. 1B–C)**. Follow-up tests in naive C57BL/6 mice showed elevated AITC responsiveness in artemin-exposed JNC neurons compared to vehicle-treated controls **(Fig. 5H–J)**. Collectively, these data suggest that alveolar macrophages, through pollutant-driven production of artemin, sensitize nociceptor neurons and thereby potentiate allergic airway inflammation. This pathway underscores a critical link between environmental pollutants and the neurogenic exacerbation of allergic responses **(SF. 3)**.

## DISCUSSION

Our research, in line with other studies, indicates that nociceptor neurons promote regulatory immunity in the context of bacterial^57^, viral^58^, or fungal^59^ infections, as well as malignancies^11^. To our knowledge, however, the impact of neuro-immunity on neutrophilic asthma is less understood. Using a model of pollution-exacerbated airway inflammation^34^, we found that ablation or silencing of nociceptor neurons prevented the induction of neutrophilic airway inflammation, revealing a potentially novel therapeutic avenue for treating refractory asthma. This result parallels our earlier findings that nociceptor neuron silencing via charged blockers of voltage-gated sodium channels ^16,22,35^, or calcium channels ^36^ can halt eosinophilic airway inflammation.

During bacterial infections, Pinho-Ribeiro et al. demonstrated that nociceptor neurons inhibit neutrophil influx and antimicrobial activity by releasing the neuropeptide CGRP^60^. In addition, Ugolini and colleagues showed that ablating Na_V_1.8^+^ nociceptor neurons during HSV-1 infection increases skin neutrophil infiltration, cytokine production, and lesion severity ^58^. Building on these findings, we used intravital imaging to study neuro–immune interactions in our model, we observed that the total number of alveolar macrophages remained unchanged in OVA-exposed mice lacking nociceptor neurons, but their displacement was significantly reduced. In contrast, pollution-exposed asthmatic mice lacking nociceptor neurons exhibited a marked decrease in neutrophil recruitment and a corresponding increase in their displacement. These shifts in cellularity occurred regardless of whether nociceptor neurons were genetically ablated (TRPV1^+^ or Na_V_1.8^+^ populations) or pharmacologically silenced with QX-314, as confirmed by flow cytometry. Although individual neutrophil behaviors—such as adherence, crawling, patrolling, and tethering—did not change, neuronal ablation effectively “removed the brakes” on neutrophil motility. This likely led to reduced neutrophil accumulation in the lung and overall dampening of lung inflammation in pollution-exacerbated asthma. These findings are consistent with the pioneering work by Baral et al., who showed that vagal TRPV1^+^ afferents suppress neutrophil recruitment and modulate lung γδ T cell numbers via CGRP signaling, thus supporting essential antibacterial clearance^4^. Related studies in influenza-infected mice revealed that vagal nociceptor deficiency significantly alters the lung immune landscape—expanding neutrophils and monocyte-derived macrophages ^5^—and transcriptional analyses further indicated disrupted interferon signaling as well as imbalances among neutrophil subpopulations in nociceptor-ablated mice. Collectively, these results highlight the bidirectional crosstalk between recruited and tissue-resident phagocytes and nociceptors in air-pollution–exacerbated allergic airway inflammation.

The aryl hydrocarbon receptor (**AhR**) is a crucial modulator of inflammation, as seen in psoriasis models, where its activation reduces inflammation, whereas its absence exacerbates disease^61,62^. Besides being expressed in alveolar macrophages, AhR is also present in ILC2s, eosinophils, and neurons. A recent preprint suggests AhR has a dual role in neural protection and axon regeneration: AhR activation in dorsal root ganglion neurons inhibits axon growth^63^. At the same time, its deletion lessens inflammation and stress signaling, driving pro-regenerative pathways^63^. Additional work indicates that AhR regulates the gut-brain axis^64^. Although we have not directly examined AhR in nociceptor neurons, our findings suggest that neurons, much like alveolar macrophages, may sense fine particulate matter via AhR. This possibility could explain the direct neuronal transcriptomic reprogramming we observed after pollutant exposure. Ongoing studies aim to probe this hypothesis using nociceptor-specific AhR knockout models.

A growing body of evidence indicates that airway-innervating sensory neurons undergo significant transcriptomic alterations in neutrophilic asthma. Although some lung innervation arises from the dorsal root ganglia (DRG)—especially for nociception in the upper thoracic segments (T1–T5)—the majority (70–80%, sometimes up to 90%) of afferent fibers originate from the JNC ^65,66^. Because the JNC mediates critical mechanosensory and chemosensory functions, our analysis focuses on this primary vagal pathway. While single-cell RNA sequencing has revealed that JNC neurons are highly heterogeneous and control breathing and tracheal reflexes ^67–72^, the molecular changes that occur during chronic inflammation remain poorly characterized.

Using an Na_V_1.8 reporter mice and intranasal retrograde tracing, we identified a subset of nociceptors innervating the lungs that are reprogrammed in allergic airway inflammation and by the asthma-driving cytokine IL-13^24^. Our transcriptomic data using sorted TRPV1^+^ neurons confirm the upregulation of *Npy1r* during allergic airway inflammation^24^ and highlight the upregulation of *Cacna1c*^73^ and *Dmxl2*^74^ during pollution-exacerbated asthma, suggesting heightened vesicle trafficking and changes in synaptic release dynamics that may influence neuronal sensitivity and plasticity in asthmatic conditions. Other genes such as *Oprm1*^75^ (mu-opioid receptor) and *Hint1*^76^ (an opioid signaling modulator) suggest altered pain or irritant sensation in the lung, potentially shaping cough reflexes or bronchospasm. Changes in cytoskeletal regulators, including *Map1b*^77^ and *Apc*^78^, indicate that JNC axons may remodel their structure and synaptic architecture, potentially altering how airway signals are relayed to central circuits. *Lifr*^79–82^ (Leukemia inhibitory factor receptor alpha), which supports neuron survival and differentiation, may also be important in inflammatory or injury contexts in the lung.

Pathway analysis reveals gene clusters involved in synaptic structure, cytoskeletal organization, and epigenetic regulation. Enrichment in terms such as “synapse”, “presynaptic active zone,” and “postsynaptic density” indicates that both neurotransmitter release and postsynaptic excitability are being recalibrated, potentially amplifying neuro-immune feedback loops and leading to maladaptive reflexes like excessive bronchoconstriction or persistent cough in neutrophilic asthma. In parallel, pathways related to “cellular component organization,” “negative regulation of inclusion body assembly,” and “protein quality control” show that neurons may upregulate protective or adaptive mechanisms in response to chronic airway inflammation. These transcriptomic alterations point to a dynamic neuro-immune crosstalk in the inflamed lung, with neurons adapting their excitability and structural integrity in response to cytokines, environmental pollutants, and inflammatory cells, potentially exacerbating or perpetuating the pathophysiology of neutrophilic asthma.

Alveolar macrophages act as a critical early warning system in the lung^45^, detecting pollutants such as FPM and initiating protective reflexes by secreting artemin. This cytokine activates and sensitizes nociceptor neurons—particularly TRPA1-expressing fibers—to noxious stimuli. Single-cell RNA sequencing identified a subset of TRPA1⁺, SSTR2⁺ neurons that express the glial cell line-derived neurotrophic factor receptor GFRα3 and are thereby sensitized by artemin. This finding aligns with earlier data showing that keratinocyte-derived TSLP can sensitize skin-innervating nociceptor neurons, promoting itch and atopic dermatitis^83^. In the lungs, germline TRPA1 knockout reduces allergic airway inflammation^84,85^, and clinical data from Genentech suggest that TRPA1 agonists are elevated in asthmatic human airways, contributing to both inflammation and hyperreactivity. GDC-0334, a selective TRPA1 antagonist they develop, mitigates airway inflammation, cough, and allergic responses in preclinical models and reduces pain and itch in humans^86^. These findings strongly support the central role of TRPA1 sensitization and nociceptor activation in driving asthma pathology.

Our prior work showed that nodose ganglion neurons, which are highly TRPA1⁺, exhibit heightened thermal sensitivity and outgrowth responses to brain-derived neurotrophic factor (**BDNF**) but do not respond to nerve growth factor (**NGF**)^43^. Here, we demonstrate that vagal nociceptors express GFRα3 and respond to artemin, which sensitizes TRPA1 channels and promotes subsequent airway inflammation. Indeed, jugular-nodose-complex (**JNC**) neurons undergo substantial transcriptomic reprogramming when exposed to OVA plus FPM, mirroring the effects of pro-asthmatic cytokines. Parallel data from other disease models support this link between inflammation-induced sensitization and elevated artemin. For example, increased *Artn* and EGR1 levels in atopic dermatitis correlate with enhanced nerve density and scratching, both of which are attenuated in EGR1-deficient mice^87^. Similarly, artemin overexpression boosts sensory neuron thermal responses, underscoring its importance in driving neuronal outgrowth and sensitivity in atopic conditions^49^. These observations raise the possibility that blocking GFRα3 may offer a novel strategy to prevent maladaptive nociceptor involvement in pollution-exacerbated asthma.

Our previous work showed that vagal nociceptor neurons can detect allergen-IgE immune complexes and respond by releasing substance P (**SP**) and vasoactive intestinal peptide (**VIP**), but not CGRP^20^. These findings were corroborated by analyzing bronchoalveolar lavage fluid (**BALF**) from asthmatic mice²⁴. Moreover, we demonstrated that SP drives mucus metaplasia in asthmatic lungs^18^ and influences antibody class switching in B cells^22^, a result that other groups have replicated^23,39,88,89^. We also observed that TRPV1⁺ nociceptor ablation impairs antigen trafficking to lymph nodes and reduces IgG production ^23,90,91^, possibly decreasing the severity or onset of allergic responses. Emerging work highlights the varied immunomodulatory roles of different neuropeptides. For instance, CGRP impairs dendritic cell migration in psoriasis ^9,92^, whereas SP enhances dendritic cell migration to lymph nodes in atopic dermatitis^8^. VIP and neuromedin U (**NMU**) boost pro-asthmatic cytokines from lung ILC2s^16,93–96^, while CGRP can have similar^97,98^ or opposing effects^99^. In lung infections, CGRP reduces neutrophil and γδ T cell infiltration and protects against *Staphylococcus aureus* pneumonia^4,100^. Conversely, CGRP exacerbates psoriasis by inducing dendritic cells to release IL-23, which activates IL–17–producing γδ T cells, intensifying inflammation^59^. Likewise, during *C. albicans* infection, sensory neurons release CGRP that stimulates IL-23 production in CD301b⁺ dendritic cells, triggering T_h_17 and γδ T cell responses (marked by IL-17A and IL-22), enhancing host defense^7^. In our model, nociceptor neuron ablation diminishes γδ T cell activation, which we posit is chiefly driven by SP/VIP, given the absence of heightened CGRP release in our asthma models^16,22^. Future work will investigate whether neurons directly regulate γδ T cells and neutrophils through these or other neuropeptides.

Previous reports show that TRPV1⁺ neuron activation elicits a local type 17 immune response that enhances host defense against *Candida albicans*^59,101^ and *Staphylococcus aureus*^7^. While our mixed OVA-FPM was expected to induce a strong T_h_17/T_h_2 component, we did not observe an increase in IL-17A in our cytokine bead array. Similarly, increases in classic asthma-driving cytokines were modest (IL-4, IL-5) or absent (IL-13). One possible explanation is that these experiments were conducted in C57BL/6 mice, rather than BALB/c mice^34^. We also noted an increase in TNFα and artemin, both of which were reduced by nociceptor neuron silencing. While we found that TNFα can reprogram nociceptor neurons^24^, we propose that this TNFα may originate from activated neutrophils^102,103^, consistent with the reduced TNFα and neutrophil activation seen in neuron-ablated mice. It is worth noting that the PM₂₅ used in this study—Standard Reference Material (SRM) 2786—was obtained from an air intake system in the Czech Republic and that we cannot rule out that it may contain trace levels of LPS that could contribute to this neutrophil–TNFα hypothesis. Our rationale for choosing SRM2786 was that it is commercially available and represents a broad spectrum of ambient air pollutants, in contrast to more specialized sources such as diesel exhaust particles. Future studies will need to validate these findings using wildfire-collected particles^104^, LPS-free air intake particles, and replication in BALB/c mice.

In summary, our study highlights how alveolar macrophages, via AhR-dependent pollutant detection and artemin secretion, sensitize TRPA1⁺ nociceptor neurons, thereby amplifying neutrophilic asthma. We propose several potential therapeutic strategies to curb neutrophilic airway inflammation: (i) targeting AhR signaling in alveolar macrophages, (ii) blocking Artemin’s interaction with GFRα3, (iii) using TRPA1 antagonists such as GDC-0334, and (iv) silencing nociceptor neurons with charged lidocaine derivatives. Collectively, these approaches offer new directions to manage the increasingly prevalent yet treatment-resistant, neutrophilic variant of asthma.

**Supplementary Figure 1.**
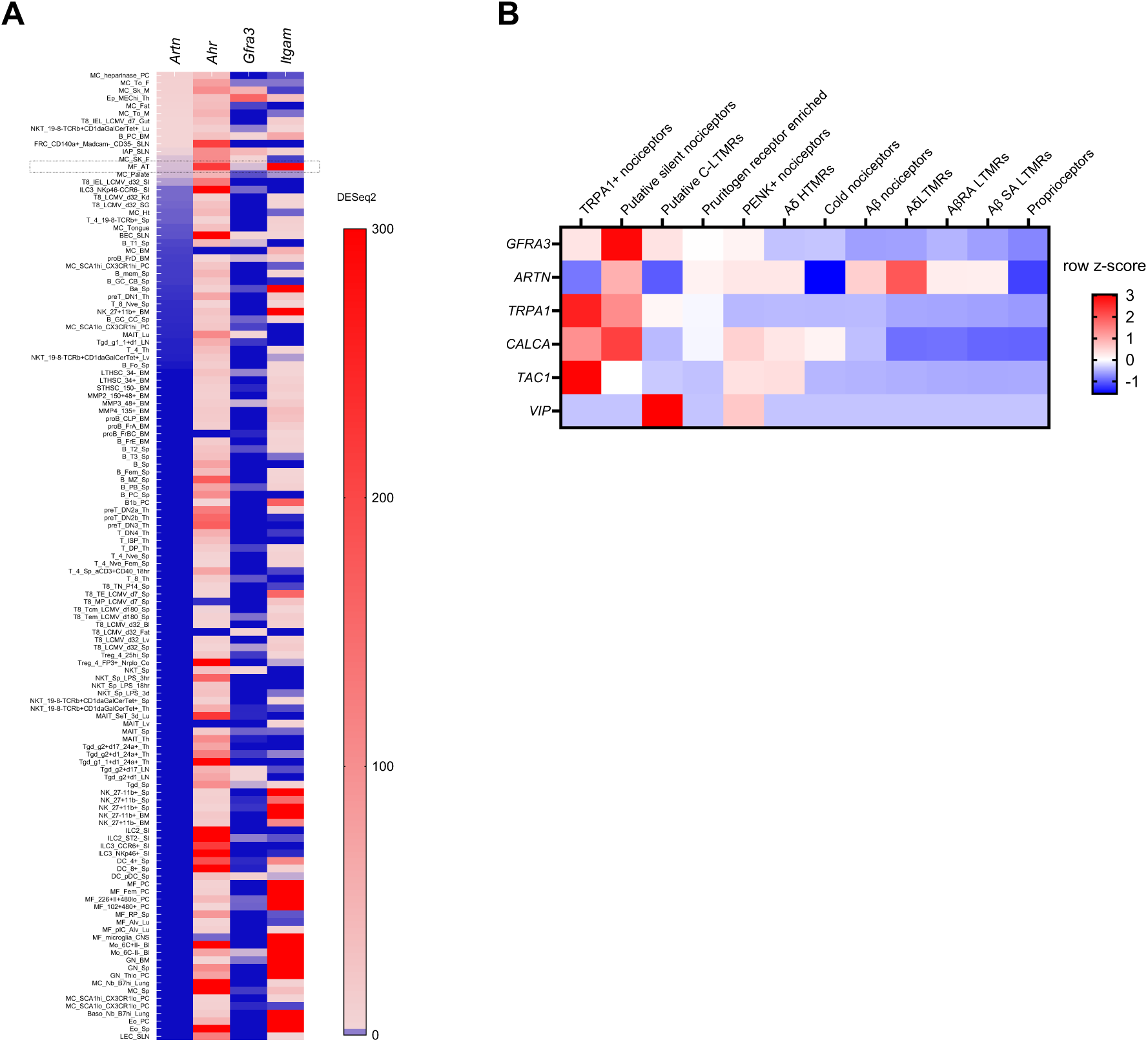
In silico analysis of *Artn* expression in mouse immune cells. **(A)** *In-silico* re-analysis of *Artn* expression in mouse immune cells using the ImmGen database^53^. *Artn* and *Ahr* are expressed in *Itgam*^+^ macrophages. Data are presented as per-gene z-scores of normalized gene expression, calculated by the median of ratios method. **(B)** *In-silico* re-analysis of the single-cell RNA-seq dataset from Tavares-Ferreira et al., ^105^ (Sensoryomics; dbGaP accession phs001158) shows that *Gfra3* is expressed in *Trpa1*-positive nociceptors, C-LTMRs, and silent nociceptors within the human dorsal root ganglion. Expression levels are reported as per-gene z-scores calculated with the median-of-ratios normalization method.

**Supplementary Figure 2.**
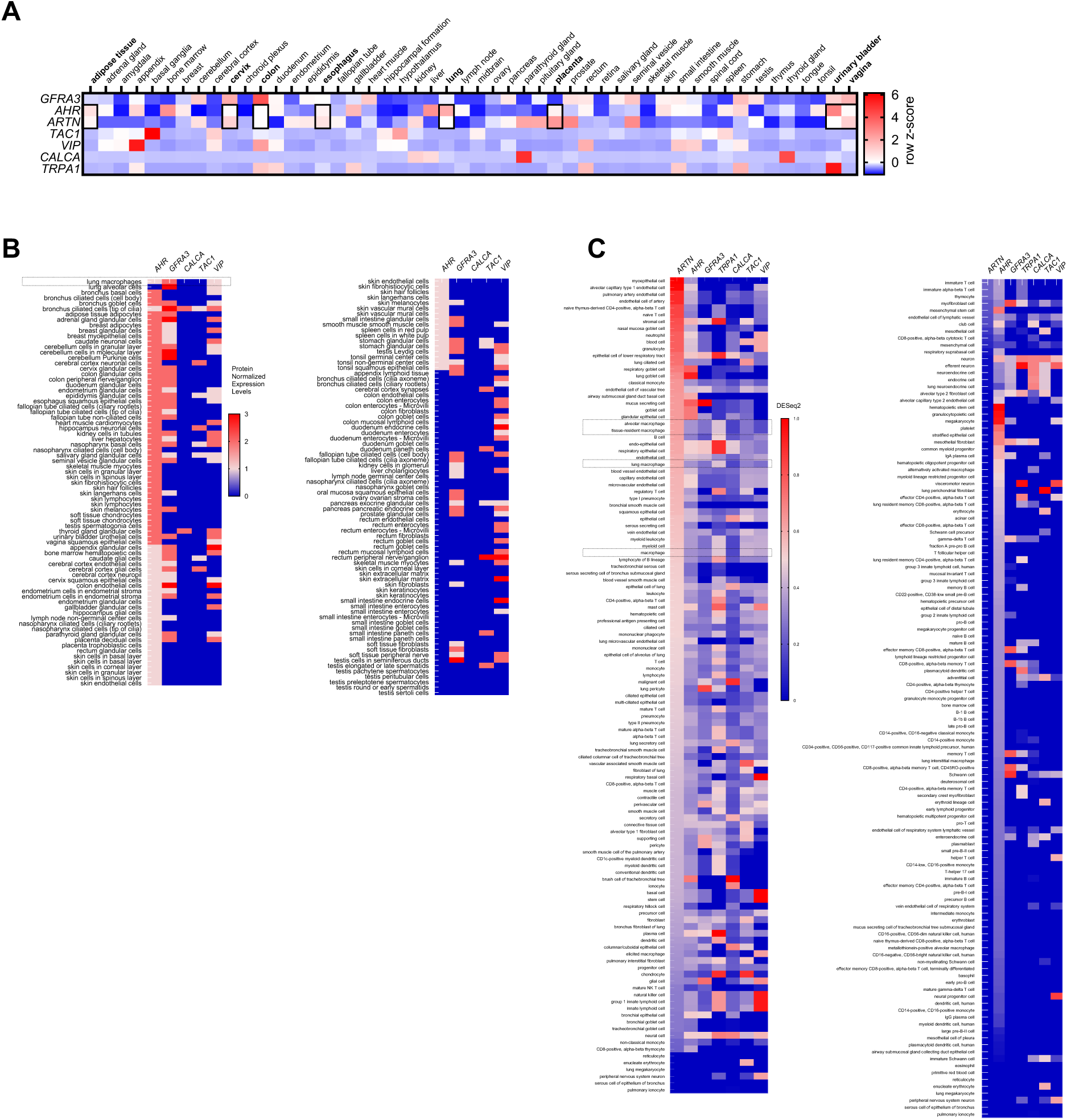
*In-silico* re-analysis of Ahr and Artn expression in human tissues. **(A)** *In-silico* re-analysis of data from Karlsson et al., ^106^ (Human Protein Atlas, proteinatlas.org) indicates that *Ahr* and *Artn* are expressed in human lung. Expression values are reported as per-gene z-scores using the median-of-ratios normalization method. Experimental details and cell clustering are described by Karlsson et al., ^106^ **(B)** *In-silico* re-analysis of data from Uhlén et al. ^55^ (Human Protein Atlas, proteinatlas.org) confirms *Ahr* protein expression in human lung macrophages, as shown by immunohistochemistry. These data are presented as protein-normalized expression levels. Experimental details and cell clustering are described by Uhlén et al. ^55^. **(C)** *In-silico* re-analysis of data from Abdulla et al. ^56^ (CELLxGENE, CZI Single-Cell Biology) reveals co-expression of *Artn* and *Ahr* in lung and alveolar macrophages from patients. Expression values are provided as per-gene z-scores, calculated by the median-of-ratios normalization method. Experimental details and cell clustering are described by Abdulla et al. ^56^.

**Supplementary Figure 3.**
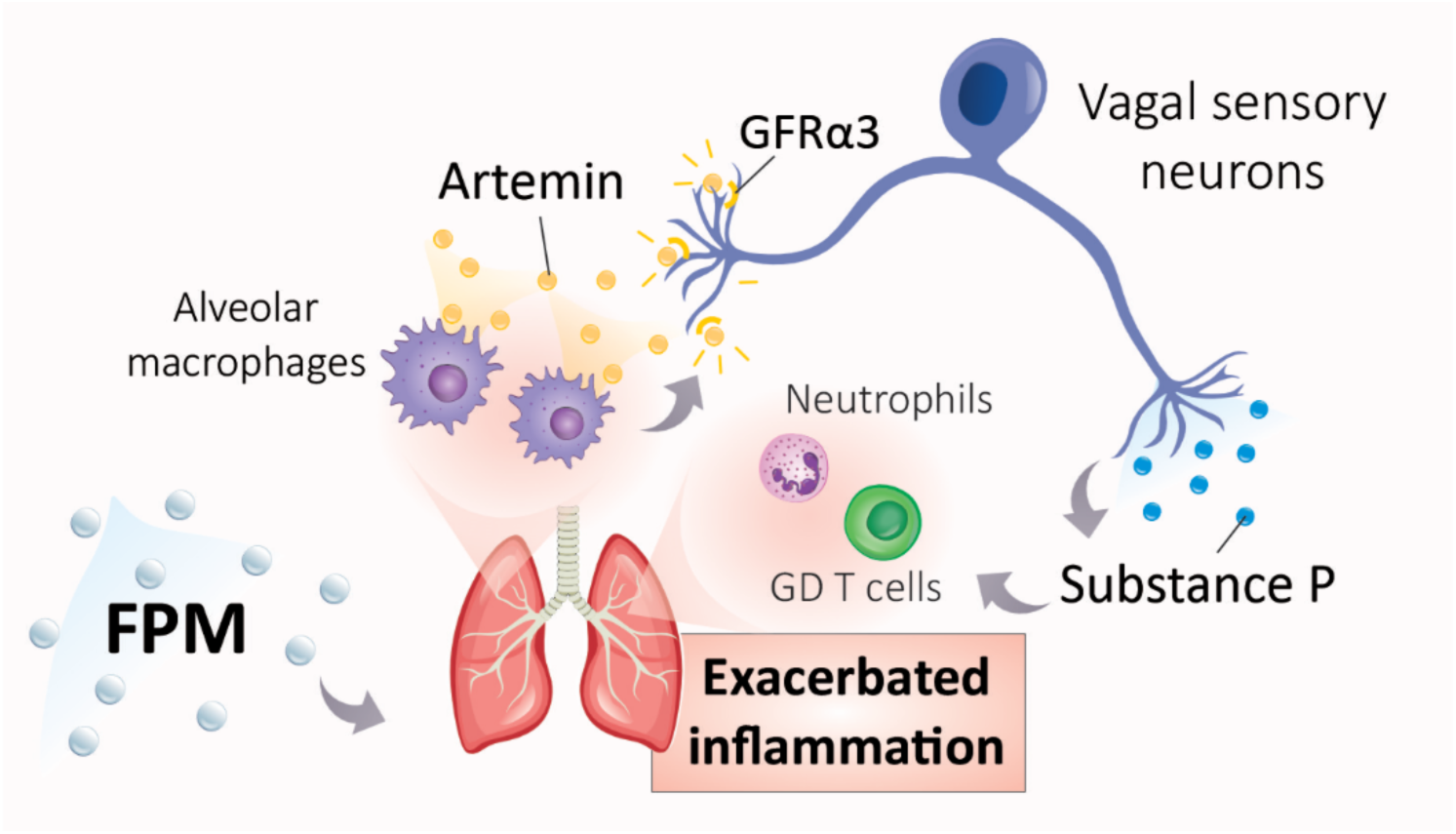
Schematic of nociceptor involvement in pollution-exacerbated allergic asthma. In our study, mice were exposed to PM₂₅ particles and ovalbumin (OVA) to model pollution-exacerbated asthma. Compared to mice exposed to OVA alone, co-exposure to PM₂₅ and OVA significantly increased bronchoalveolar lavage fluid (BALF) neutrophils and lung γδ T cells levels. To counteract this heightened airway inflammation, we administered intranasal QX-314—a charged lidocaine derivative—at the peak of inflammation, effectively normalizing BALF neutrophil levels. Ablation of TRPV1⁺ nociceptor neurons produced a similar effect. Further analysis with calcium imaging revealed that neurons from the jugular-nodose complex in pollution-exposed asthmatic mice were more sensitive via their TRPA1 channels. Levels of TNFα and the growth factor artemin were also elevated in the BALF of these mice, returning to normal following nociceptor ablation. We identified alveolar macrophages as the source of artemin, which they secrete upon sensing fine particulate matter (FPM) through aryl hydrocarbon receptors. Artemin, in turn, heightened TRPA1 responsiveness to its agonist (mustard oil), thereby exacerbating airway inflammation. Our findings suggest that silencing nociceptor neurons can disrupt this pathway, offering a novel therapeutic approach to mitigate neutrophilic airway inflammation driven by pollution.

**Supplementary Table 1.** Differentially expressed genes and pathway analysis of vagal nociceptors in pollution-exacerbated asthma.

Naïve 6–10 weeks male and female TRPV1^cre^::tdTomato^fl/wt^ mice were either subjected to a pollution-exacerbated asthma protocol, to the classic ovalbumin (OVA) protocol or remained naïve. On day 17, jugular-nodose complex (JNC) neurons were harvested and dissociated, and TRPV1⁺ (tdTomato⁺) neurons were FACS-purified to remove stromal and non-peptidergic cells before being processed for RNA sequencing. The different tabs show the DESeq2 identified for each of these conditions. Other tabs show GO terms enriched in each condition, analyzed using the web-based tool g:Profiler.

**Supplementary Video 1.** Intravital recording of alveolar macrophage motility.

6–10 weeks old male and female littermate control (Na_V_1.8^wt^::DTA^fl/wt^ denoted as NaV1.8^WT^) and nociceptor-ablated (Na_V_1.8^cre^::DTA^fl/wt^ denoted as Na_V_1.8^DTA^) mice were sensitized via intraperitoneal injection of an emulsion containing ovalbumin (OVA; 200 µg/dose) and aluminum hydroxide (1 mg/dose) on days 0 and 7. On day 10, phagocytes were labeled by intranasal injection of PKH26 (25 pmol/dose). Mice were then challenged intranasally with OVA (50 µg/dose) alone or in combination with fine particulate matter (FPM; 20 µg/dose) on days 14–16. Alveolar macrophage intravital imaging was performed on day 17 and is presented as a 1-hour time-lapse video.

**Supplementary Video 2.** Intravital recording of neutrophil motility.

Male and female littermate control (Na_V_1.8^wt^::DTA^fl/wt^ denoted as Na_V_1.8^WT^) and nociceptor-ablated (Na_V_1.8^cre^::DTA^fl/wt^ denoted as Na_V_1.8^DTA^) mice (6–10 weeks old) were sensitized via an intraperitoneal injection of an ovalbumin (OVA; 200 µg/dose) and aluminum hydroxide (1 mg/dose) emulsion on days 0 and 7. On days 14–16, mice were challenged intranasally with OVA (50 µg/dose) alone or in combination with fine particulate matter (FPM; 20 µg/dose). Immediately before intravital imaging on day 17, an intravenous Ly6G antibody was administered to label neutrophils. The resulting recording is presented as a 20-minute time-lapse video.

## MATERIALS AND METHODS

### Animals

All procedures involving animals adhered to the guidelines of the Canadian Council on Animal Care (CCAC), McGill University (xxx) and the Queen’s University Animal Care Committee (UACC, protocol 2384). Mice were housed in individually ventilated cages with free access to water and food under 12-hour light cycles.

Parental strains C57BL/6 (# 000664), DTA^fl/fl^ (# 010527, # 009669), tdTomato^fl/fl^ (# 007914), TRPV1^cre/cre^ (# 017769) and Na_V_1.8^cre/cre^ (# 036564) were purchased from The Jackson Laboratory. Male and female mice were bred in-house and used at 6–12 weeks of age. Crosses were performed to generate the following genotypes: Na_V_1.8^cre/wt^::DTA^fl/wt^, TRPV1^cre/wt^::DTA^fl/wt^, TRPV1^cre/wt^::tdTomato^fl/wt^, and littermate control, namely: Na_V_1.8^wt/wt^::DTA^fl/wt^, TRPV1^wt/wt^::DTA^fl/wt^. Male and female progeny mice were used between 8 and 16 weeks of age.

### Ovalbumin Model of Allergic Airway Inflammation

On days 0 and 7, mice were sensitized by intraperitoneal (i.p.) injections of 200 μL of a solution containing 1 mg/mL grade V ovalbumin (OVA; Sigma, #A5503) and 5 mg/mL aluminum hydroxide (Sigma, #239186) in phosphate-buffered saline (PBS, ThermoFisher, #10010023). On days 14–16, mice were anesthetized with isoflurane (2.5%; CDMV, #108737) and intranasally instilled daily with 50 μg OVA in 50 μL PBS with or without fine particulate matter (FPM; 20 µg/dose; NIST 2786). Control mice were sensitized but did not undergo challenges. Unless otherwise indicated, mice were sacrificed on day 17.

### Neuron silencing

In some experiments, QX-314 (Tocris, #1014; 5 nmol in 50 µL) or PBS was administered intranasally on day 16 to control asthmatic mice (as detailed in the asthma protocol). The mice were euthanized on day 17, and bronchoalveolar lavage fluid, lung tissues, and the jugular-nodose complex were collected.

### Bronchoalveolar lavage

Bronchoalveolar lavage fluid was harvested in anesthetized mice, following a tracheal incision, by lavaging twice with 1 ml of either PBS or FACS buffer (2% FBS and 1 mM EDTA in PBS) through a Surflo ETFE IV Catheter 20G × 1″ (Terumo Medical Products, # SR-OX2025CA). The lavage fluid was centrifuged at 350 × g for 6.5 minutes, and the supernatant was collected for ELISA analysis. The cell pellet was resuspended, subjected to RBC lysis (Cytek, # TNB-4300-L100 or Gibco, # A1049201), and stained for surface markers before flow cytometry.

### Flow cytometry

Single-cell suspensions derived from bronchoalveolar lavage fluid or lung samples were stained with Ghost Dye Violet 510 (Cytek, # 13-0870-T100) and appropriate antibody cocktails in PBS. Cells were incubated at 4°C for 30 minutes, then fixed with 10% neutral buffered formalin (Sigma Aldrich, # HT501128) at room temperature for 15 minutes before data acquisition. To assess eosinophil and neutrophil infiltration in BALF, cells were stained with fluorochrome-conjugated antibodies against CD45 (clone: 30-F11), CD90.2 (clone: 53-2.1), CD11b (clone: M1/70), CD11c (clone: N418), Ly6C (clone: HK1.4), Ly6G (clone: 1A8), and Siglec-F (clone: 1RNM44N). For γδ T cell analysis in lung tissue, staining included CD45 (clone: 30-F11), TCRγδ (clone: GL3), CD90.2 (clone: 53-2.1), and lineage markers TCRβ (clone: H57-597), CD19 (clone: 1D3/CD19), NK1.1 (clone: PK136), CD11b (clone: M1/70), CD11c (clone: N418), F4/80 (clone: BM8), and FcεRIα (clone: MAR-1), obtained from Biolegend or Thermo Fisher Scientific. Data were acquired using a BD FACS Canto II system.

### Lung tissue harvesting

After diaphragm incision and transcardial perfusion with 10 ml of PBS, lung tissues were dissected, minced with razor blades, and either placed in TRIzol™ Reagent (Invitrogen, # 15596026) for RNA extraction or transferred into a digestion buffer consisting of 1.6 mg/ml collagenase type 4 (Worthington LS004189) and 100 µg/ml DNase I (Roche, # 11284932001) in supplemented DMEM^107^. The tissues were digested for 45 minutes at 37°C with mechanical dissociation through 18-gauge needles after 30 minutes, followed by filtration through a 70 µm nylon mesh and RBC lysis. Cells were resuspended in FACS buffer for flow cytometry or fluorescence-activated cell sorting or in FBS-supplemented DMEM for in vitro stimulation in 96-well plates at 37°C with 5% CO₂, after which supernatants were collected.

### Alveolar macrophage culture

Alveolar macrophages were obtained from the BALF of naïve mice, where they represented approximately 95% of the recovered cells. After centrifugation and RBC lysis, these cells were seeded at 3 × 10^5^ per well in 96-well plates containing DMEM (Gibco 11965092) supplemented with 1 mM sodium pyruvate (Gibco, # 11360070), 2 mM GlutaMAX (Gibco, # 35050061), 100 U/mL penicillin and 100 µg/mL streptomycin (Corning, # 30-002-CI), 10 mM HEPES (Gibco, # 15630080), and 10% FB Essence (VWR 10805-184), and cultured overnight. They were then stimulated with 100 µg/ml FPM (NIST, # 2786) for 1–4 hours, followed by RNA extraction for quantitative PCR analysis.

### Neuron culture

The jugular-nodose complex (JNC) was collected from anesthetized mice following exsanguination and placed in a digestion buffer containing 1 mg/ml (325 U/ml) collagenase type 4 (Worthington, # LS004189), 2 mg/ml (1.8 U/ml) Dispase II (Sigma, # 04942078001), and 250 µg/ml (735.25 U/ml) DNase I (Roche, # 11284932001) prepared in supplemented DMEM without FB Essence. This mixture was incubated at 37°C for 60 minutes to ensure enzymatic digestion, followed by mechanical dissociation through progressive pipetting with tips of decreasing diameter and final passage through a 25-gauge needle. The cell suspension underwent density gradient centrifugation at 200 × g for 20 minutes at low acceleration and deceleration, layering 150 mg/ml bovine serum albumin (BSA; Hyclone, # SH30574.02) in PBS to separate the cells. The bottom fraction was collected, RBC lysed and seeded onto glass-bottom dishes (Abidi, # 81218) coated with 50 µg/ml laminin (Sigma, # L2020) and 100 µg/ml poly-D-lysine (Sigma, # P6407). Cells were cultured overnight in Neurobasal-A medium (Gibco, # 10888022) supplemented with 1 mM sodium pyruvate (Gibco, # 11360070), 2 mM GlutaMAX™ (Gibco, # 35050061), 100 U/mL penicillin, 100 µg/mL streptomycin (Corning, # 30-002-CI), 10 mM HEPES (Gibco, # 15630080), B-27 supplement (Gibco 17504-044), 50 ng/ml mouse nerve growth factor (NGF; Gibco, # 13257-019), 2 ng/ml mouse glial-derived neurotrophic factor (GDNF; Novus, # NBP2-61336), and cytosine-β-D-arabinofuranose (Thermo Scientific, # J6567106). In some experiments, 100 ng/ml artemin or 50 µM HCl (as vehicle control) were added in vitro in place of NGF or GDNF. This culture system was subsequently used for calcium imaging.

### Real-time quantitative PCR (qPCR)

qPCR was performed on stimulated alveolar macrophages that were lysed using TRIzol Reagent and stored at –80°C until RNA extraction. RNA from sorted cells was extracted using the PureLink RNA Micro Scale Kit (ThermoFisher, # 12183016). In contrast, RNA from lung tissues or lung cell suspensions was extracted with the E.Z.N.A.® Total RNA Kit I (Omega Bio-tek, # R6834). Extraction procedures followed the manufacturers’ instructions, including phenol-chloroform purification and mixing with an equal volume of isopropanol. Complementary DNA (cDNA) was synthesized using the SuperScript VILO Master Mix (Invitrogen, # 11755050), with 1–2 µg of RNA as the template in each reaction. Quantitative PCR was carried out with PowerUp SYBR Green Master Mix (Applied Biosystems, # A25742), using 50–100 ng of cDNA and 200 nM of each primer. The reactions were run on either a Mic qPCR Cycler (Bio Molecular Systems) or a CFX Opus Real-Time PCR System (Bio-Rad Laboratories). The primer pair for *Artn* was Forward: 5′-TGATCCACTTGAGCTTCGGG-3′ and Reverse: 5′-CTCCATACCAAAGGGGTCCTG-3′.

### Calcium imaging recording

Cultured neurons were loaded with 5 µM fura-2 AM (Cayman Chemical Company, # 34993) and incubated at 37°C for 40 minutes. After incubation, the cells were washed four times with standard external solution (SES; Boston BioProducts, # C-3030F) and maintained in this solution during imaging. The Fura-2 signals were recorded and used for downstream analyses. Agonists, diluted in SES, were delivered using a ValveLink 8.2 system (AutomateScientific) equipped with 250 µm Perfusion Pencil® tips (Automate Scientific) and controlled by Macro Recorder (Barbells Media, Germany). Between drug injections, SES flow was maintained to ensure a complete washout of each agonist. Imaging for Fura-2 experiments was conducted with a NIKON ECLIPSE Ti2 Inverted Microscope using an S Plan Fluor ELWD 20X objective lens to optimize UV light transmission. Images were captured every 3 or 4 seconds using sCMOS cameras such as PCO.Edge 4.2 LT (Excelitas Technologies), Prime 95B (Teledyne Photometrics), or Orca Flash 4.0 v2 (Hamamatsu Photonics). Regions of interest were manually delineated in NIS-Elements (Nikon), and the F340/F380 ratios were exported to Excel (Microsoft 365) for further analysis^108^. The data were compressed by calculating a maximum value every 15 seconds for all subsequent evaluations.

### Intravital microscopy

6–10 weeks old male and female littermate control (Na_V_1.8^wt^::DTA^fl/wt^ denoted as Na_V_1.8^WT^) and nociceptor-ablated (Na_V_1.8^cre^::DTA^fl/wt^ denoted as Na_V_1.8^DTA^) mice were sensitized via intraperitoneal injection of an emulsion containing ovalbumin (OVA; 200 µg/dose) and aluminum hydroxide (1 mg/dose) on days 0 and 7. On day 10, phagocytes were labeled by intranasal injection of PKH26 Red Fluorescent Cell Linker Kit (Sigma, # PKH26PCL) at 25 pmol/dose in Diluent B. Mice were then challenged intranasally with OVA (50 µg/dose) alone or in combination with fine particulate matter (FPM; 20 µg/dose) on days 14–16. Alveolar macrophage intravital imaging was performed on day 17 and is presented as a 1-hour time-lapse video.

Lung images were acquired using a Nikon CSU-X1 multichannel spinning-disk confocal upright microscope with a protocol adapted from previously published methods^44,45^. Mice were anesthetized with 10 mg/kg xylazine hydrochloride and 200 mg/kg ketamine hydrochloride, and body temperature was maintained at 37°C with a heating pad (World Precision Instruments). The right jugular vein was cannulated for additional anesthetic as needed and for injecting anti-mouse Ly6G Alexa Fluor 647 (BioLegend, clone: 1A8, # 127610) to label neutrophils. After exposing the trachea and inserting a catheter connected to a small rodent ventilator (Harvard Apparatus), the mouse was placed in a right lateral decubitus position. A small incision was made between ribs 4 and 5 to create an opening of about 1.5 cm, and an intercostal lung window was carefully fitted and stabilized by a vacuum of roughly 20 mmHg. Time-lapse images were acquired without delay using a 20X water-immersion objective (numerical aperture 1).

Images are presented as maximum-intensity projections of z-stacks. Alveolar macrophage movement was tracked for over 1 hour, and neutrophil behavior was recorded for 20 minutes. Cell displacement was quantified using the ICY software’s manual tracking plugin. Neutrophil behavior was classified into adherent, tethering, crawling, or patrolling using Imaris (Oxford Instruments) spot tracking. Track Duration and Track Speed Mean were used to define each behavior per field of view (FOV). Tethering was determined by a Track Duration under 150 seconds for cells that rapidly entered the FOV before arresting and exiting again. Adherent cells remained immobile for more than 150 seconds, with a Track Speed Mean of ≤ 0.03 µm/s. Crawling cells were motile with steady movement that persisted for at least half of the video’s duration; these had a Track Speed Mean > 0.03 µm/s and a Track Duration > 600 seconds. Patrolling cells shared the rapid entry observed with tethering cells but, instead of briefly arresting continued crawling and exited the FOV, showing a Track Speed Mean > 0.03 µm/s and a Track Duration > 150 seconds but < 600 seconds.

### Bulk RNA sequencing

TRPV1^cre/wt^::tdTomato^fl/wt^ mice were sensitized via intraperitoneal injection with a mix of grade V OVA (200 µg/dose; Sigma-Aldrich A5503) and Imject® Alum (1 mg/dose; ThermoFisher 77161) on days 0 and 7. Subsequently, they underwent intranasal challenges with OVA (50 µg/dose), with or without fine particulate matter (FPM; 20 µg/dose; NIST 2786), from day 14 to 16. Control mice were sensitized but not challenged. The mice were euthanized on day 17, jugular-nodose complex (JNC) were collected, and tdTomato^+^ cells from naive, OVA-challenged, and OVA-FPM co-exposed mice were sorted by FACS ^24^, and total RNA was extracted following established protocols. Library preparation was carried out at the Institut de Recherche en Cancérologie et en Immunologie (IRIC) of the Université de Montréal. RNA quality was assessed using an Agilent Bioanalyzer, ensuring a minimum RNA Integrity Number (RIN) of 7.5. Libraries were prepared using a poly(A)-enrichment, single-stranded RNA-seq strategy (KapaBiosystems, KAPA RNA Hyperprep Kit, #KR1352) and sequenced on an Illumina NextSeq500 platform with 75-cycle single-end reads.

Basecalling was performed using Illumina RTA 2.4.11, and demultiplexing was conducted with bcl2fastq 2.20, allowing for one mismatch in the index. Trimmomatic was used to remove adapter sequences and low-quality bases from the 3′ end of each read. The remaining high-quality reads were aligned to the GRCm38 mouse genome using STAR v2.5.11, which also generated gene-level read counts. Differential expression analysis was performed using DESeq2 on these read counts, normalized by the DESeq2 pipeline. Log2 fold changes and –log10 p-values were calculated from the normalized data, and genes were considered differentially expressed if their adjusted p-value (false discovery rate, FDR) was below 0.05. Further data analysis and visualization were conducted in RStudio. Bulk RNA-sequencing raw and processed data have been deposited in the NCBI’s gene expression omnibus (GSE298583).

### Enzyme-linked immunosorbent assay (ELISA)

ELISA was used to measure artemin levels in BALF (R&D Systems, # DY1085-05) following the manufacturer’s instructions. Inflammatory cytokines in BALF were detected with a Cytometric Bead Array Flex Set from BD Biosciences: Master Buffer Set (# 558266), IL-1β (# 560232), IL-4 (# 558298), IL-5 (# 558302), IL-6 (# 558301), IL-10 (# 558300), IL-13 (# 558349), IL-17A (# 560283), IFNγ (# 558296), MCP-1 (# 558342), and TNF (# 558299), also used according to the manufacturer’s guidelines.

### In-silico analysis of mouse immune cells expression profile using the Immgen database

Using the publicly available Immgen database^53^ we proceed to an in-silico analysis of RNA-sequencing data (DESeq2 data) of various mouse immune cells. As per Immgen protocol, RNA-sequencing reads were aligned to the mouse genome GENCODE GRCm38/mm10 primary assembly (GenBank assembly accession GCA_000001635.2) and gene annotations vM16 with STAR 2.5.4a. The ribosomal RNA gene annotations were removed from general transfer format file. The gene-level quantification was calculated by featureCounts. Raw reads count tables were normalized by median of ratios method with DESeq2 package from Bioconductor and then converted to GCT and CLS format. Samples with less than 1 million uniquely mapped reads were automatically excluded from normalization. Experimental details are defined in www.immgen.org/Protocols/ImmGenULI_RNAseq_methods.pdf.

### In silico analysis of RNA-Seq data

In silico analysis of RNA-Seq data involved extracting information from Kupari et al.’s supplementary materials (GSE124312)^51^, with clusters based on that publication’s designations. Additional data from Zhao et al., (GSE192987)^52^ were reanalyzed using R and plotted with UMAP to visualize the co-expression of relevant genes. Bulk JNC sequencing datasets were analyzed with DESeq2.

### In-silico analysis of lung cancer patients’ tumor expression profile using single-cell RNA sequencing

We performed an *in-silico* analysis of single-cell RNA-sequencing data from mouse lung tumor–infiltrating CD45⁺ cells in non–small cell lung cancer (NSCLC) models (GSE127465)^109^, accessed via the publicly available Broad Institute Single-Cell Portal. The expression levels of *Ahr, Artn, Calca, Vip, Tac1,* and *Trpa1* were plotted within CD45⁺ myeloid cell populations using RStudio. Additionally, we conducted a Spearman correlation analysis to explore relationships among these genes. Gene expression data are presented as normalized counts per ten thousand. We extracted the relevant expression values and used RStudio to generate dot plots for individual cells in each myeloid population, as well as to perform correlation analyses of the selected genes. Experimental and clustering details can be found at Zilionis et al., ^109^.

### In-silico analysis of human lung-tissue gene expression using the Human Protein Atlas (HPA) Single Cell Type Atlas

We retrieved the lung dataset generated by Karlsson et al. ^106^ from the HPA Single Cell Type Atlas, which pools single-cell RNA-sequencing reads into annotated clusters and reports expression as per-gene z-scores after median-of-ratios normalization. Using R (v4.3.2), we downloaded the cluster-level expression matrix, isolated all clusters and extracted transcript values for *Gfra3, Ahr, Artn, Calca, Tac1, Vip* and *Trpa1*. The positive z-scores observed in the “lung” clusters confirm that Ahr mRNA is expressed in patient lung tissue. Because the Karlsson pipeline equalizes library depth before z-score transformation, these values permit direct comparison of Ahr abundance to the whole-body baseline captured by the same atlas. Additional, experimental and clustering details can be found at Karlsson et al., ^106^.

### In-silico analysis of AHR protein abundance in human lung macrophages using Human Protein Atlas immunohistochemistry data

We retrieved the antibody-based tissue microarray summary from Uhlén et al. ^55^ via the Human Protein Atlas pathology portal, which lists per-tissue immunoreactivity for each antibody as categorical scores (“not detected,” “low,” “medium,” “high”). From the raw TSV file, we report protein expression (*Ahr, Artn, Gfra3, Calca, Vip, Tac1*) across cell types and tissue types. Because the HPA pipeline reports these scores after internal normalization of DAB signal across replicate tissue cores, the categorical value can be compared directly across tissues without further scaling. This processed result confirms constitutive AHR protein presence in lung macrophages in situ. Additional, experimental and clustering details can be found at Uhlén et al. ^55^.

### In-silico co-expression analysis of AHR and ARTN in human lung macrophages using the CZ CELLxGENE Discover single-cell atlas

We accessed the lung single-cell RNA-seq data curated by Abdulla et al. ^56^ through the CZ CELLxGENE Discover platform and downloaded the cell-level expression matrix together with standardized cell-type annotations. Raw UMI counts were normalized to ln (CPTT + 1) values during ingestion into the atlas, a transformation that equalizes library depth and stabilizes variance while preserving the full gene–cell count matrix for downstream analyses. Additional, experimental and clustering details can be found at Abdulla et al. ^56^.

### G-Profiler and Go-term

The top 50 differentially expressed genes (based on DESeq p-values) from each comparison were submitted to the web-based tool g:Profiler ^110^ for enrichment analysis (g:GOSt). The resulting pathway enrichments—derived from multiple databases, including Gene Ontologies (GO; covering Molecular Function, Biological Process, and Cellular Component sub-ontologies) ^111^, the Kyoto Encyclopedia of Genes and Genomes (KEGG) ^112^, Reactome (REAC) ^113^, WikiPathways (WP) ^114^, TRANSFAC (TF) ^115^, miRTarBase (MIRNA), the Comprehensive Resource of Mammalian Protein Complexes (CORUM) ^116^, and the Human Phenotype Ontology (HP) ^117^—are listed in Supplementary Table 1.

### Data availability

Bulk RNA-sequencing raw and processed data have been deposited in the NCBI’s gene expression omnibus (GSE298583). Processed data can also be accessed in the supplementary Table 1. Additional information and raw data are available from the lead contact upon reasonable request.

### Statistics

P values ≤ 0.05 were considered statistically significant. One-way ANOVA, and Student’s t-tests were conducted using GraphPad Prism, while DESeq2 and Seurat analyses, including their statistical tests, were performed in RStudio.

### Replicates

The number of replicates (n) for each experiment is specified in the figure legends and represents the number of animals for in vivo data. For in vitro experiments, replicates may be culture wells or dishes, animals, fields of view during microscopy, or individual neurons in calcium imaging. All experiments included different preparations from distinct animals to ensure biological reproducibility.

### Exclusion

*One replicate in the naïve mice group of the RNA-sequencing analysis (*Figure 2*) was excluded because the PCA indicated it as an outlier. No other data were excluded from the study*.

## ACKNOWLEDGEMENTS.

ST’s work is supported by the Canadian Institutes of Health Research (193741, 407016, 461274, and 461275), the Canadian Foundation for Innovation (44135), the Canadian Cancer Society Emerging Scholar Research Grant (708096), the Knut and Alice Wallenberg Foundation (KAW 2021.0141, KAW 2022.0327), the Swedish Research Council (2022-01661), the Natural Sciences and Engineering Research Council of Canada (RGPIN-2019-06824), and the NIH/NIDCR (R01DE032712). AT’s work is supported by the Canadian Institute of Health Research (186176). Salary support for JCW was provided by the Fonds de recherche du Québec – Santé, the Canadian Allergy, Asthma, and Immunology Foundation, Asthma Canada, and CIHR. Salary support for AK was provided by the Canada Graduate Scholarship Masters.

